# Genomic variance of the 2019-nCoV coronavirus

**DOI:** 10.1101/2020.02.02.931162

**Authors:** Carmine Ceraolo, Federico M. Giorgi

## Abstract

There is rising global concern for the recently emerged novel Coronavirus (2019-nCov). Full genomic sequences have been released by the worldwide scientific community in the last few weeks in order to understand the evolutionary origin and molecular characteristics of this virus. Taking advantage of all the genomic information currently available, we constructed a phylogenetic tree including also representatives of other *coronaviridae*, such as Bat coronavirus (BCoV) and SARS. We confirm high sequence similarity (>99%) between all sequenced 2019-nCoVs genomes available, with the closest BCoV sequence sharing 96.2% sequence identity, confirming the notion of a zoonotic origin of 2019-nCoV. Despite the low heterogeneity of the 2019-nCoV genomes, we could identify at least two hyper-variable genomic hotspots, one of which is responsible for a Serine/Leucine variation in the viral ORF8-encoded protein. Finally, we perform a full proteomic comparison with other *coronaviridae*, identifying key aminoacidic differences to be considered for antiviral strategies deriving from previous anti-coronavirus approaches.

## Introduction

*Coronaviridae* (CoVs) are the largest known single stranded RNA viruses [1]. They have been categorized in three groups, based on phylogenetic analyses and antigenic criteria [2], specifically: 1) alpha-CoVs, responsible for gastrointestinal disorders in human, dogs, pigs and cats; 2) beta-CoVs, including the Bat coronavirus (BCoV), the human Severe Acute Respiratory Syndrome (SARS) virus and the Middle Eastern Respiratory Syndrome (MERS) virus; 3) gamma-CoVs, which infect avian species.

Very recently, a novel beta-CoVs Coronavirus (2019-nCoV) originating from the province of Wuhan, China, has been causally linked to severe respiratory infections in humans. At the time of writing, 14,441 cases of 2019-nCoV-related pneumonia cases have been reported in China, plus 118 cases from 23 other countries. There are currently 315 deaths linked to this pathogen (source: World Health Organization report, 02-Feb-2020). Phylogenetic relationships between Bat and Human *coronaviridae* have been discovered for SARS [3] and more recently also for 2019-nCoV [4], suggesting events of inter-species transmissions [5].

No vaccine for 2019-nCoV has been publicly released, but a World effort has arisen towards the characterization of the molecular determinants and evolutionary features of this novel virus. An initial comparison of 10 genomic sequences from 2019-nCoV specimens has reported a low heterogeneity of this viruses with intersample sequence identity above 99.9% [6]. There are currently 54 2019-nCoV complete genome sequences from the Global Initiative for Sharing all Influenza Data (Gisaid [7]) and from GenBank [8], plus two partial sequences obtained by the Spallanzani hospital in Rome, Italy (also from Gisaid).

In this short report, we set out to characterize the heterogeneity of all 2019-nCoV genomes and proteomes available at the moment of the study, comparing them to other representative *coronaviridae*,specifically SARS, MERS and BCoV. We will generate phylogenetic trees of the 2019-nCoV cases and apply entropy-based analyses of position-wise variance and a Categorical Principal Component analysis as an alternative method to estimate the sequence distance between all analyzed viruses.

## Results

### Phylogenetic Analysis

We collected 53 full genomic 2019-nCoV sequences from the Gisaid database (Supplementary Table S1), plus the GenBank-deposited sequence from the Wuhan seafood market pneumonia virus isolate Wuhan-Hu-1 (NC_045512.2) and two partial sequences from Italian isolates (EPI_ISL_406959 and EPI_ISL_406960). To compare 2019-nCoVs with closely related viral species, we obtained six sequences from distinct human SARS genomes from GenBank (the reference NC_004718.3, plus the genomes AY274119.3, GU553363.1, DQ182595.1, AY297028.1 and AY515512.1). We also obtained six Bat coronavirus genomic sequences (DQ022305.2, DQ648857.1, JX993987.1, KJ473816.1, MG772934.1, EPI_ISL_402131). Finally, as more distantly related beta-CoVs we analyzed also MERS genomes from GenBank entries JX869059.2 and KT368829.1.

Similarly to a previous report with 10 virus specimens [6], we detected a very high conservation between the 56 analyzed 2019-nCoV genomes, with sequence identity above 99%. We found a bat CoV genome (Gisaid EPI_ISL_402131) with 96.2% sequence identity (and query coverage above 99%) to the 2019-nCoV reference sequence (NC_045512.2), while the previously reported closest bat CoV (bat-SL-CoVZC45) has a sequence similarity of 88% [6]. The reference human SARS genome (NC_004718.3) appears more distant from the reference 2019-nCoV, with sequence identity of 80.26% and query coverage of 98%.

We aligned all the 70 coronavirus sequences using MUSCLE [9] and inferred the evolutionary relationships between these sequences with a Tamura-Nei Maximum Likelihood estimation [10] with 100 bootstraps for model robustness estimation.

The results are shown in Figure 1 as a phylogenetic tree representation. All the human 2019-nCoV appear very similar to each other, despite the different locations of sampling. Bat coronaviridae appear to be the closet homologs. Two specific specimens gathered in 2013 and 2015 in China from the bat species *Rhinolophus affinis* and *Rhinolophus sinicus* appear to be located between the Bat coronavirus and the human 2019-nCoV groups, supporting the notion of a zoonotic transfer from bats to humans [4]. Human SARS sequences group with Bat coronavirus sequences more distantly related to 2019-nCoV genomes. Finally, MERS genomes are the most genetically distinct amongst the other sequences.

**Figure 1.**
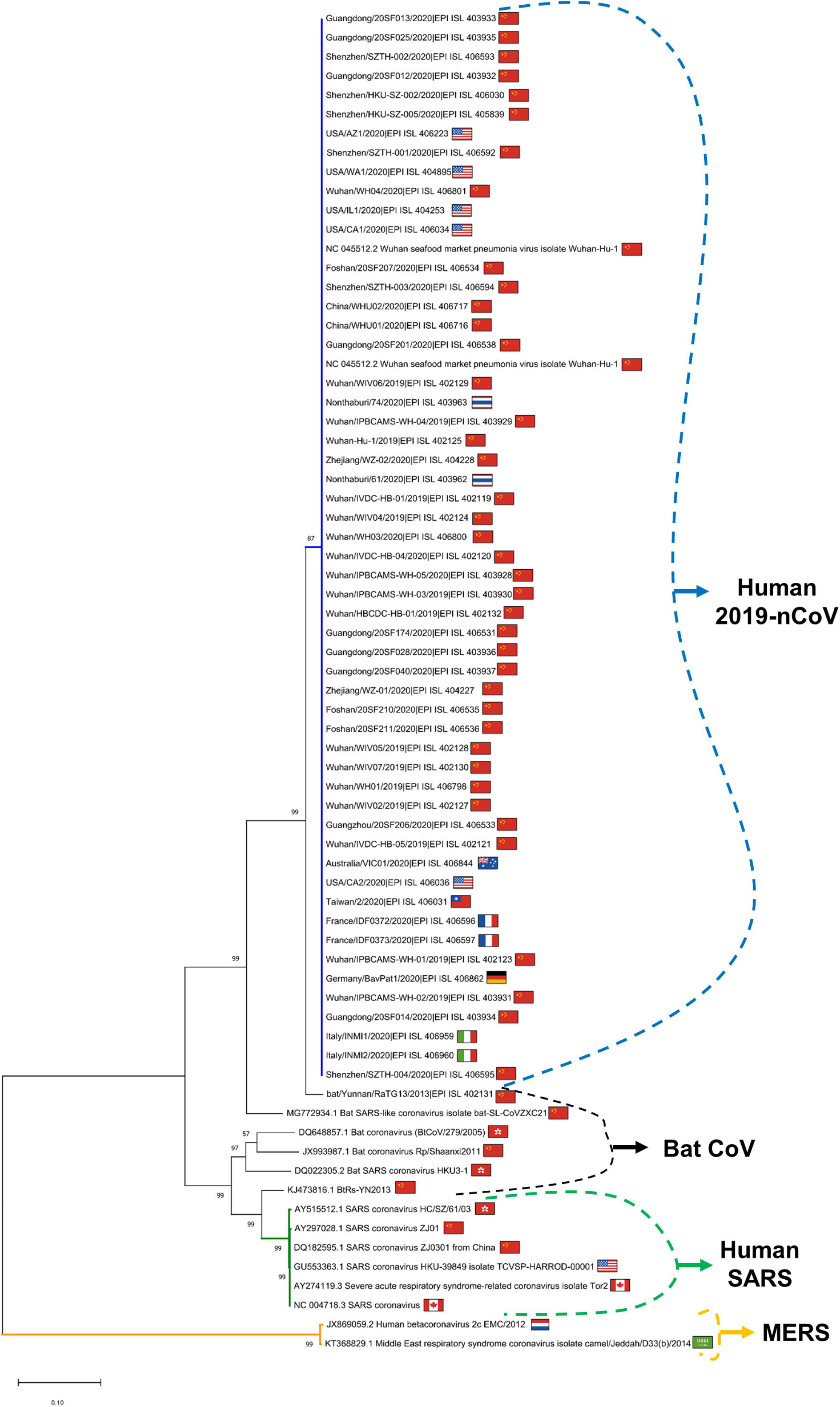
Phylogenetic tree of all the 2019-nCov sequences available at 02-Feb-2020 (branches shown in blue), plus six Bat coronavirus sequences (default black, as they are split in multiple taxa), six Human SARS (green) and 2 MERS (orange). The percentage of bootstraps supporting each branch is reported. Branches corresponding to partitions reproduced in less than 50% bootstrap replicates are collapsed.

A purely topological representation of a bootstrapped Maximum Likelihood tree (Supplementary Figure S1) shows that 2019-nCoV sequences are highly similar to each other, with poor support to the existence of distinct subgroups.

The global Multiple Sequence Alignment (MSA) is available as Supplementary File S1.

### Global Genomic Variance

Given the high homogeneity between 2019-nCoV genomes, we developed a novel method to classify genomic sequences, based on categorical Principal Component Analysis (CATPCA, [11]). Briefly, this analysis finds the eigenvectors describing the highest variance within a categorical dataset, like ours. Our dataset derived from the MUSCLE MSA of of 70 genomes and generated 32,206 positions: the categories in each coordinate could be A(Adenine), C(Cytosine), G(Guanine), (Thymine, although being a ssRNA virus, it would be more appropriate to use U - Uracil), N (Nucleotide, uncertain location: very rare in this dataset, and accounting for only 9 positions, or 0.0004% of all the data).

Our analysis shows similar results to phylogenetic tree representations. In Figure 2 A, we show the catPCA of the first components for all analyzed genomes. The MERS/nonMERS grouping accounts for the largest variance, while SARS and SARS-like BCoVs cluster together). While 2019-nCoV constitute a tightly similar cluster, the two bat virus sequences MG772934.1 and EPI_ISL_402131 appear to be linking the human 2019-nCoV to the bat *coronaviridae*.

**Figure 2.**
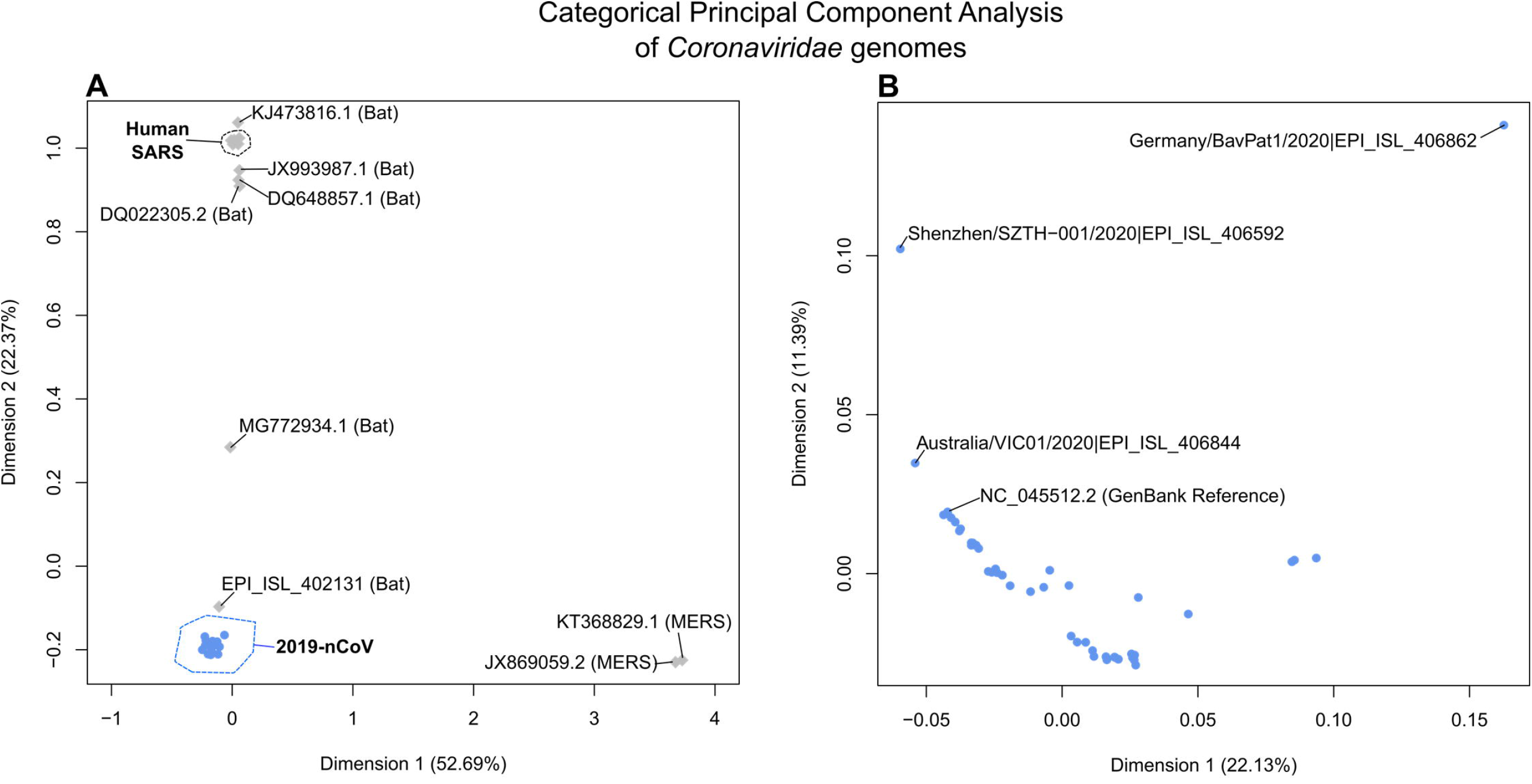
Categorical Principal Component Analysis of the projected variance in the entire *coronaviridae* genome dataset considered in this study (panel A) and in the 2019-nCoV subset only (panel B). Human 2019-nCoV are shown as blue circles, other genomes as grey diamonds.

A catPCA analysis on the sole 2019-nCoV sequences highlights some internal variability (Figure 2 B), with two likely outliers identified in the genome EPI_ISL_406862 (collected in Germany) and EPI_ISL_4O6592 (collected in Shenzhen, China).

### Genomic variance estimation within 2019-nCoV genomes

Although the variability within the 2019-nCoV genomes is very low, we set out to discover possible hot spots of hypervariability within the viral population. We analyzed the ~30,000nt of multiple genome alignments performed on the 54 full 2019-nCoV genomes. Our analysis shows that these viruses have largely the same genomic arrangement as the SARS species [12]. A large gene encoding for a polyprotein (ORF1ab) at the 5’ end of the genome is followed by 4 major structural protein-coding genes: S = Spike protein, E = Envelope protein, M = Membrane protein and N = Nucleocapsid protein. There are also at least 6 other accessory ORFs (Open Reading Frames) (Figure 3 A).

**Figure 3.**
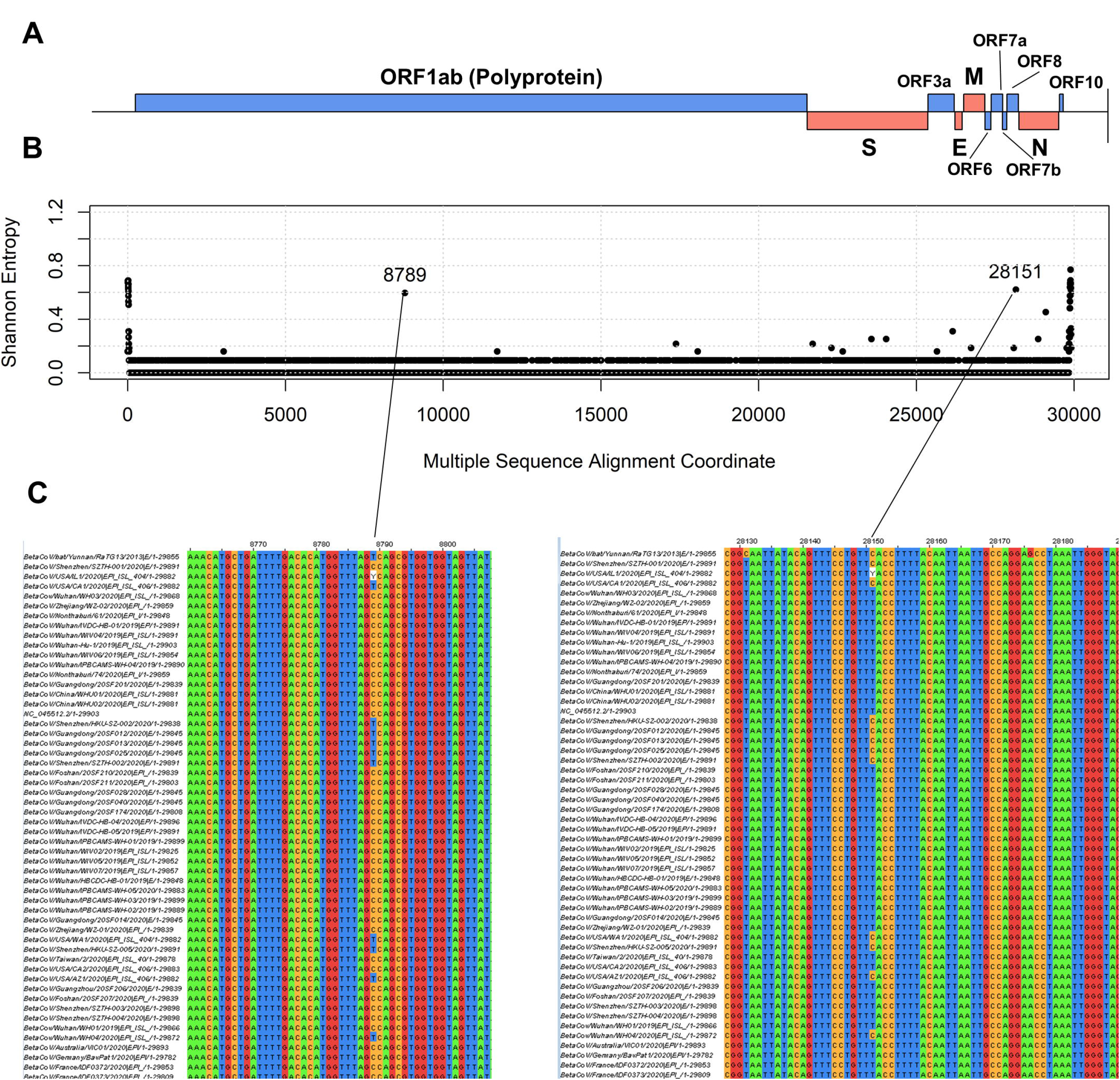
Variability within 54 2019-nCoV full genomic sequences. **(A)** location of major structural protein-encoding genes (red boxes; S = Spike protein, E = Envelope protein, M = Membrane protein, N = Nucleocapsid protein) and accessory protein ORFs (blue boxes) on the meta-genomic sequence derived from the MSA of all genomes. **(B)** Shannon entropy values across genomic locations. The two coordinates with the highest entropy (excluding the 5’ and 3’ highly variable UTRs) are indicated. **(C)** Zoom-in of the MSA describing the two most variable locations in the core genome, in the ORF1ab (left) and in ORF8 (right).

For each position of the multi-aligned 54 2019-nCoV, we calculated Shannon Entropy as a measure of the position variability [13]. Apart from the 5’ and 3’ ends, likely non-protein coding and less homogeneous, we identified two hotspots of hypervariability at positions 8789 and 28151 (Figure 3 B and C).

Position 8789 witnesses the presence of either T (U) or C, and it falls within the polyprotein gene. It causes a synonymous variation in the nucleotide triplet encoding for Serine 2839 (aminoacid coordinates based on the reference genome NC_045512.2), so it is likely not to introduce phenotypical differences between the different strains.

On the other hand, position 28151 falls within ORF8 and is characterized by the presence of either a C or a U. This causes a Ser/Leu change in aminoacid (aa) 84, which can affect the conformation of the peptide, given that Serine is a polar aminoacid, and Leucine is nonpolar. Aa84 appears to be non-conserved also across other *coronaviridae* (Figure 4 A, black arrow).

**Figure 4.**
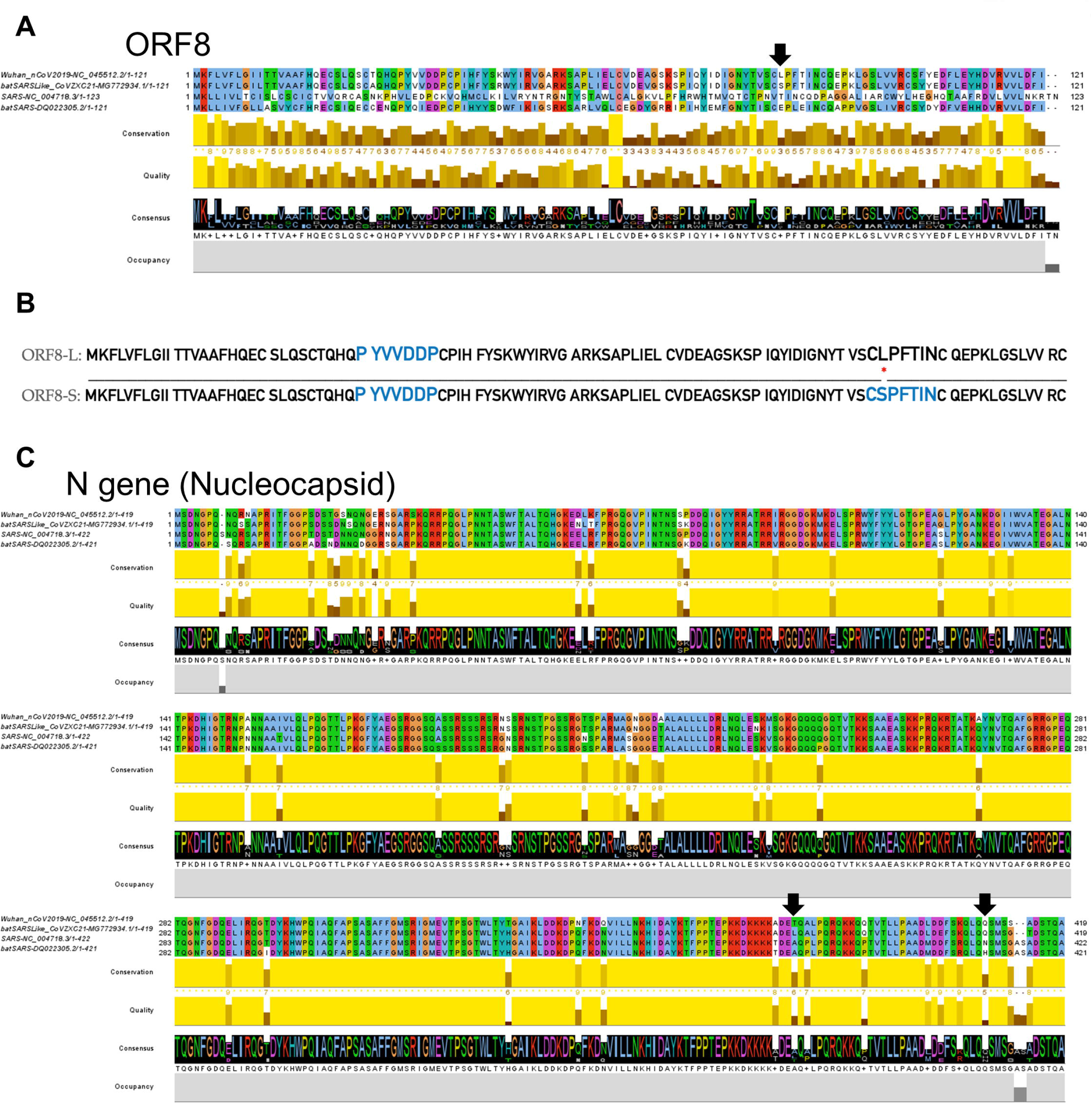
**(A)** MSA of the *coronavirus* proteins encoded by ORF8 in representative sequences for the 2019-nCoV (Wuhan), BCoVs (batSARSLike and batSARS) and SARS viruses. **(B)** Alignment of ORF8-L and ORF8-S, two isoforms varying at aminoacid position 84 (indicated by a red dot). The diagram shows in larger blue the Globplot-predicted intrinsically disordered regions in the two isoforms. **(C)** MSA of the *coronavirus* Nucleocapside proteins.

We analyzed the alternative isoforms of 2019 nCoV ORF8-aa84 alternative isoforms, dubbed ORF8-L (Leucine) and ORF8-S (Serine). Unfortunately, no crystal structures of close homologs to the ORF8 protein are available for a reliable homology modelling to measure the structural impact of this aminoacid substitution. The closest 3D model to 2019-nCoV ORF8 available in Protein Data Bank [14] is a short 22 aminoacid stretch in the protein entry 6P65, with a non-significant E-value of 0.848. We therefore employed *de novo* methods to infer structural features of ORF8. One important effect we could detect is a significant effect of Serine in ORF8-S in inducing structural disorder in the protein C-terminal portion, which is not predicted to be present in the ORF8-L (Figure 4 B), using the Russell/Linding algorithm [15]. Moreover, it did not escape our attention that the ORF8-S could theoretically generate a novel phosphorylation target for the mammalian host Serine/Threonine kinases of the host organism. So, we searched for ORF8 homologous substrates in the *Mammalia* NCBI nr protein database, but could not find matches within E-value threshold of 1.

### Protein Conservation within 2019-nCoV and between other beta-*coronaviridae*

We performed a cross-species analysis for all proteins encoded by the 2019-nCoV and its relatives. We therefore analyzed the protein sequences encoded by all ORFs in these genomes

- Wuhan NC_045512.2 (GenBank reference genome for 2019-nCoV)
- Bat Coronavirus bat-SL-CoVZXC21 MG772934.1 (bat virus similar to 2019-nCoV)
- Bat SARS coronavirus HKU3-1 DQ022305.2 (bat virus more distantly related to 2019-nCoV)
- SARS NC_004718.3 (GenBank reference genome for SARS)

Our analysis shows a close homology for all proteins with Bat sequence MG772934.1 (>80%), and more distant with the other Bat sequence and SARS reference. Query (2019-nCoV) coverage was always above 99.0%. Generally, we could observe a high conservation for structural proteins E, M and A across the beta-coronavirus family, while accessory proteins (especially ORF8) seem to have much stricter evolutionary constraints (Table 1).

**Table 1:** percentage identity between the human reference nCoV proteins and other coronavirus representatives.

| Gene | Bat MG772934.1 | Bat DQ022305.2 | SARS NC_004718.3 |
| --- | --- | --- | --- |
| ORF1ab (polyprotein) | 95.15% | 85.78% | 86.12% |
| S (Spike) | 80.32% | 76.04% | 75.96% |
| Orf3a | 92.00% | 72.99% | 72.36% |
| E (Envelope) | 100% | 94.74% | 94.74% |
| M (Membrane) | 98.65% | 90.99% | 90.54% |
| ORF6 | 93.44% | 67.21% | 68.85% |
| ORF7a | 88.43% | 88.52% | 85.25% |
| ORF7b | 93.02% | 79.07% | 81.40% |
| ORF8 | 94.21% | 57.02% | 30.16% |
| N (Nucleocapsid) | 94.27% | 89.55% | 90.52% |
| ORF10 | 73.20% | 74.23% | 72.45% |

On average, nCoV shares 91.1% of protein sequences with Bat virus MG772934.1, 79.7% with Bat virus DQ022305.2 and 77.1% with the SARS proteome.

For further visual confirmation, via MSA, of the conservation between SARS and 2019-nCoV, especially in structural proteins, please refer also to Figure 4 C and Supplementary Figures S2 and S3. As previously observed, the N protein in 2019-nCoV differs from the SARS ortholog in the structurally relevant aminoacids 380 and 410 [4].

## Discussion

Our results highlight a high level of conservation within 2019-nCoV genomes sequenced so far, and a clear origin from other beta-CoVs, specifically BCoVs, SARS and MERS. Our analysis confirms previous results highlighting the Bat coronavirus as a likely evolutionary link between the SARS viruses and the current epidemic 2019-nCoV [4]. We could confirm this result both with standard phylogenetic analysis and a newly developed visualization method for genomic distances, based on CATPCA.

The similarity between 2019-nCoV and the closest Bat relative is very high: all proteins in the coronavirus proteome (with the exception of ORF10) have identities of above 85%, with full conservation of the genome length (~30kb). We could report also the specific aminoacids that changed between SARS and nCoV, with potential implication in epitope definition and possible repurposing of anti-SARS drugs and vaccines.

Our analysis found low variability (>99% sequence identity) within the 56 2019-nCoV genomes available at the time of writing, with only two core positions of high variability, one a silent variant in the ORF1ab locus, and the other as an aminoacid polymorphism in ORF8. The mutation in ORF8 resulting in one of its two variants, ORF8-L and ORF8-S, is predicted to be affecting the structural disorder of the protein. Specifically, the amino acidic region aa83-aa89 is more likely to be disordered in the ORF8-S isoform.

In conclusion, our analysis confirms low variability within the new epidemic virus 2019-nCoV sequenced specimens, while highlighting at least two nucleotide positions of higher variability within protein coding regions, and specific aminoacid divergences compared to BCoVs and SARS [4]. These findings shed a cautiously optimistic light on the possibility of finding effective treatment for this novel coronavirus, starting from already existing *anti-beta-coronaviridae* compounds [16], which will be dealing with a relatively homogenous viral population.

## Methods

All genomic sequences were collected On 02-Feb-2020 from GenBank [8] or Gisaid [7].

MSA was performed using MUSCLE v3.8.31 [9].

MSA visualization was generated *via* Jalview v 2.11.0 [17].

Phylogenetic model generation and tree visualization was done using MEGAX v 10.1.7 [18], using the Maximum Likelihood method and Tamura-Nei model [10]. The tree structure was validated by running the analysis on 100 bootstrapped input datasets [19].

Categorical Principal Component Analysis was performed on R version 3.6.1 using the package FactoMineR [11]. Specifically, a MSA FASTA file from MUSCLE is loaded in R and converted into a categorical matrix, with genomes as rows and nucleotide coordinates as columns. Factors are defined as A, C, G, T, N or - (gap), as described in results. Then, the FactoMineR Multiple Correspondence Analysis (MCA) algorithm is run with default parameters and custom R functions are used to plot the component values for each genome.

Pairwise protein identity and coverage was calculated using BLAST protein v2.6.0 [20] with BLOSUM62 matrix and default parameters. Nucleotide sequence identity and coverage were calculated using BLAST nucleotide v2.6.0 [20].

Prediction of structural protein disorder was performed using GLOBPLOT2, an implementation of the Russell/Linding algorithm [15].

## Supplementary Materials

**Supplementary Figure S1**: topological tree for *coronaviridae* genomes.

**Supplementary Figure S2**: MSA of M, E, S, ORF3a, ORF6, ORF7a, ORF7b and ORF10 proteins in four different *coronaviridae*.

**Supplementary Figure S3**: MSA of ORF1ab proteins in four different *coronaviridae*.

**Supplementary Table S1**: annotation of Gisaid 2019-nCoV used in this study.

**Supplementary File S1**: MSA of all coronavirus genomes in this study.

## Acknowledgments

We would like to thank the Italian Ministry of University and Research for funding. Also, we would like to acknowledge the fruitful discussions with our colleagues Daniele Mercatelli, Simone Di Giacomo and Giorgio Milazzo. Also, a big acknowledgement to Eleonora Fornasari, for the help with graphics.

