## Supplementary figures and images for "Genomic variance of the 2019-nCoV coronavirus"

### Supplementary Figure S1

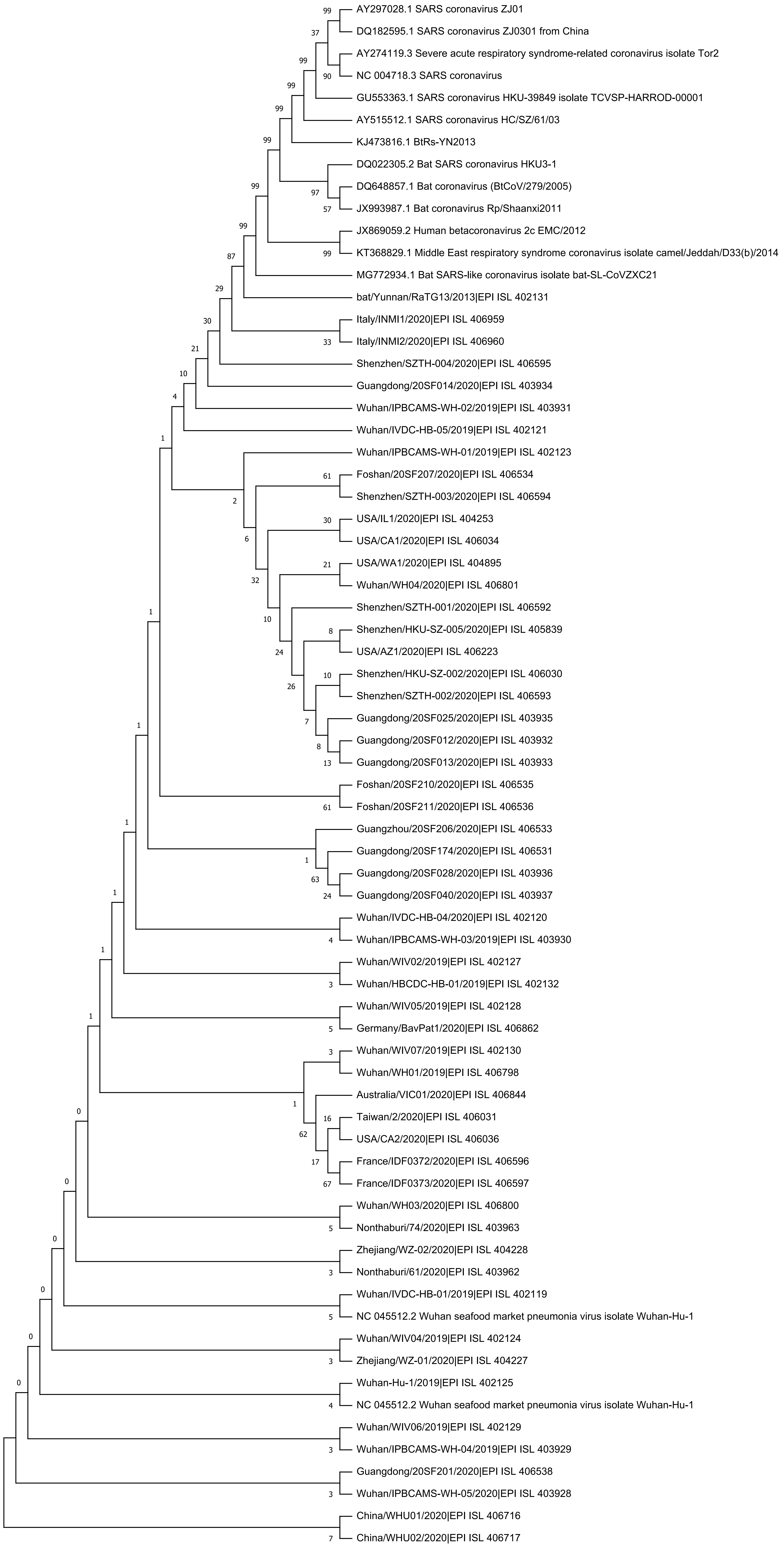

### Supplementary Figure S3

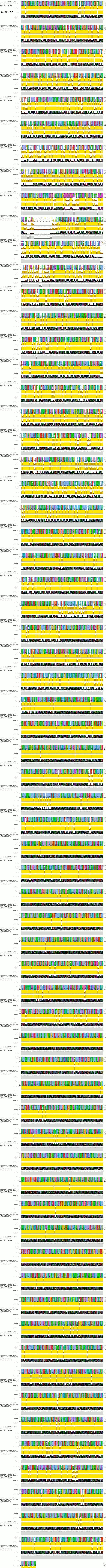
