## Supplementary Figure S2 for "Genomic variance of the 2019-nCoV coronavirus"

### M gene (Membrane)

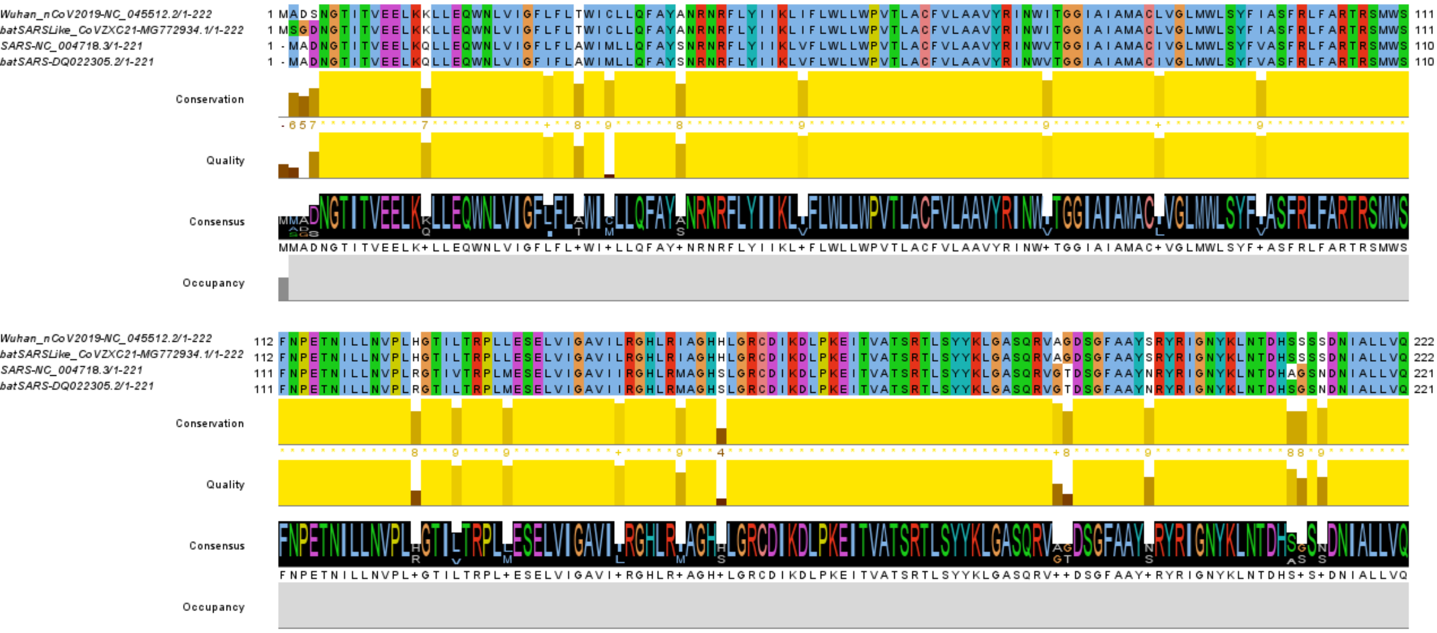


### E gene (Envelope)

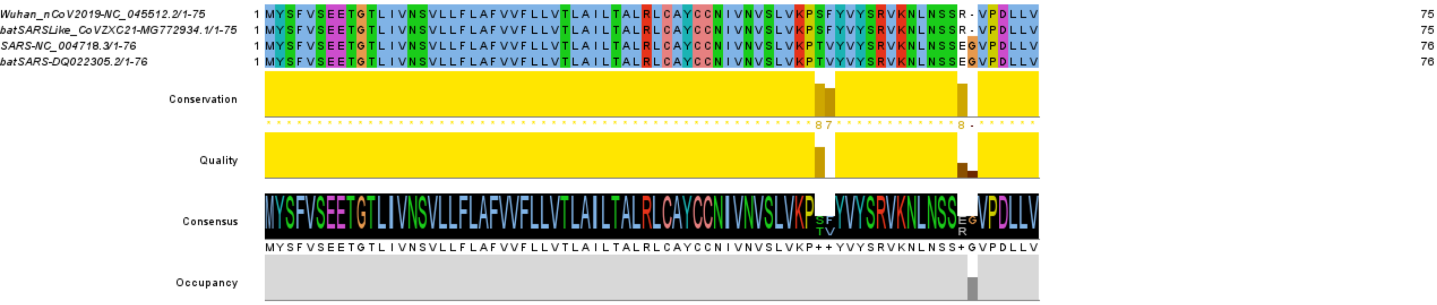


### ORF6

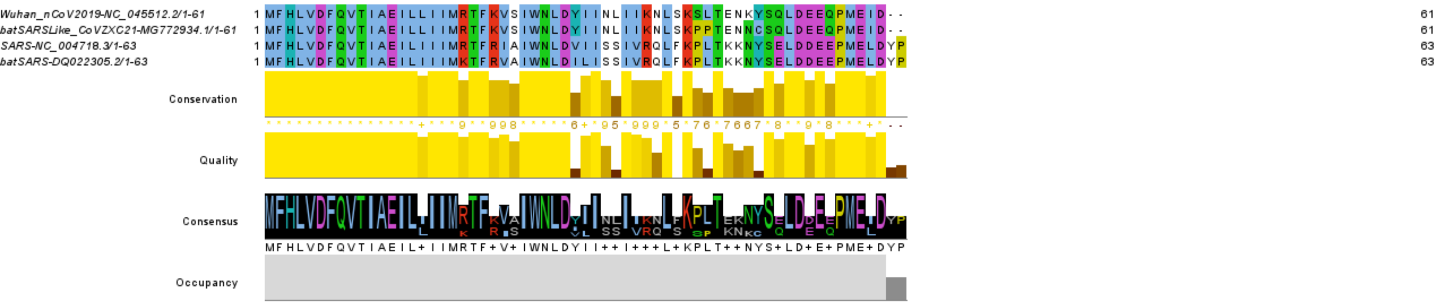


**Supplementary Figure S2A**

### ORF7a

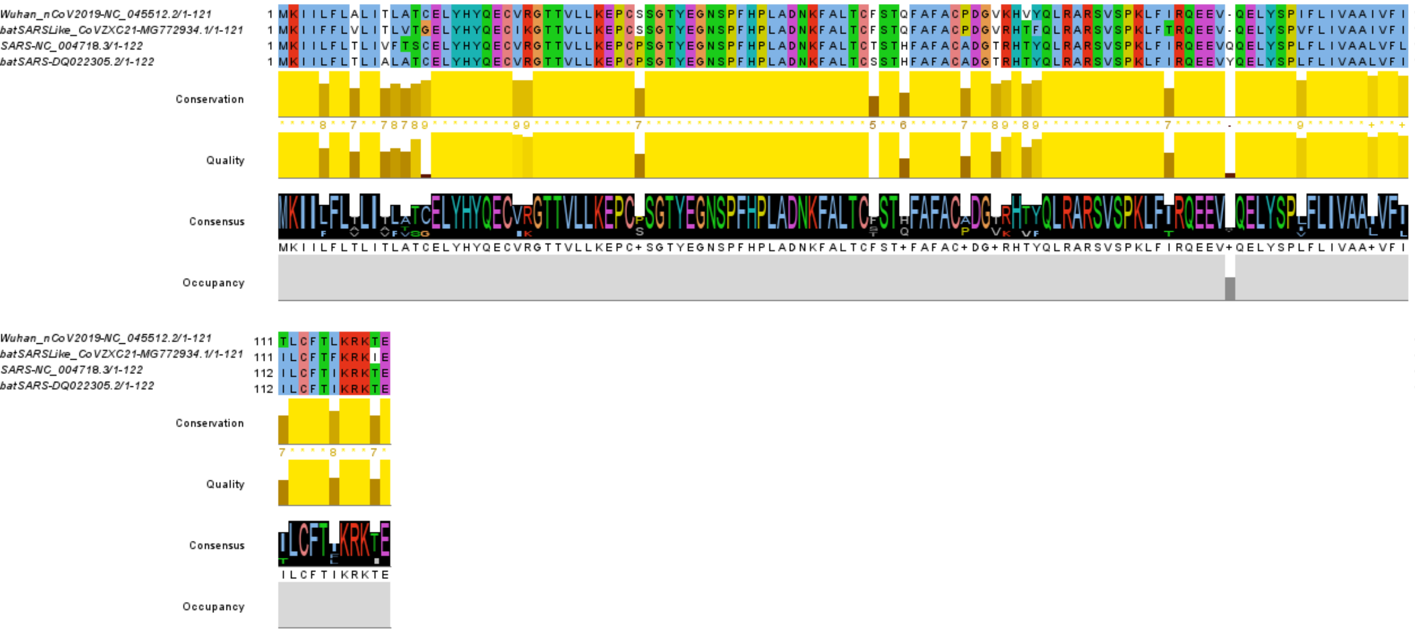


### ORF7b

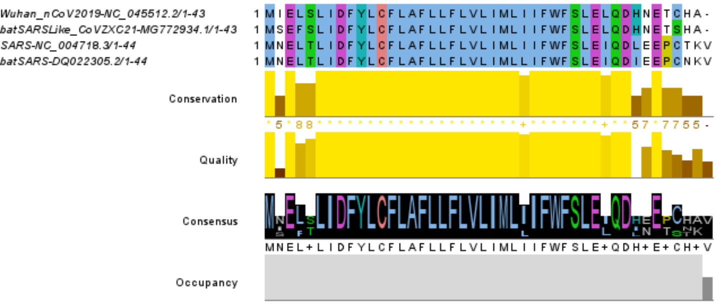


### ORF10

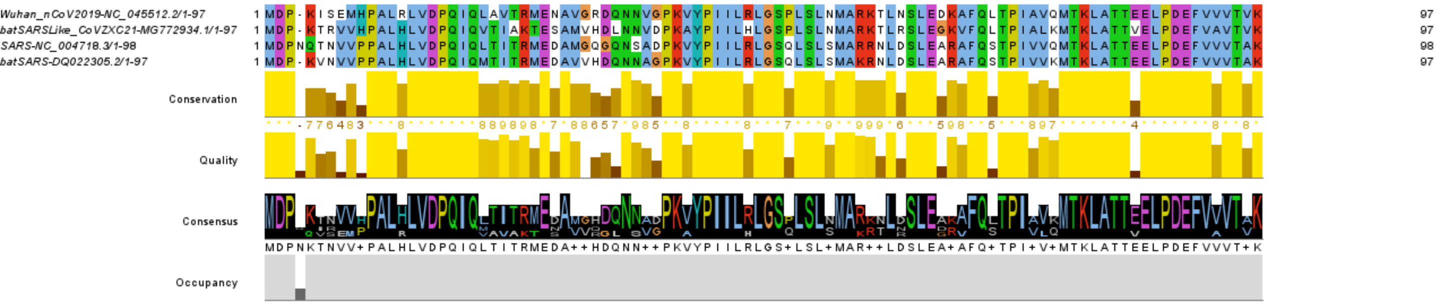


**Supplementary Figure S2B**

### ORF3a

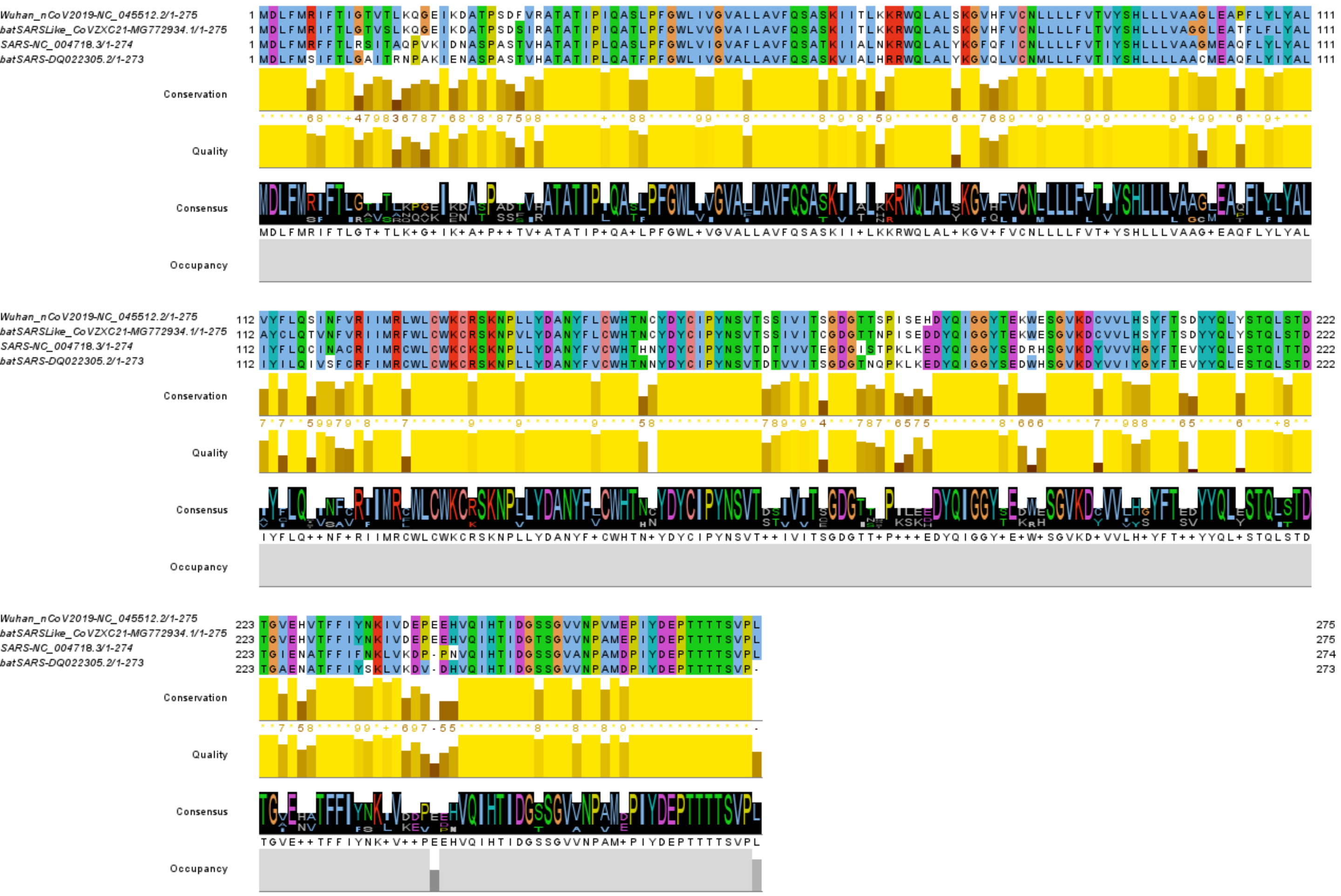


**Supplementary Figure S2C**

### S gene (Membrane)

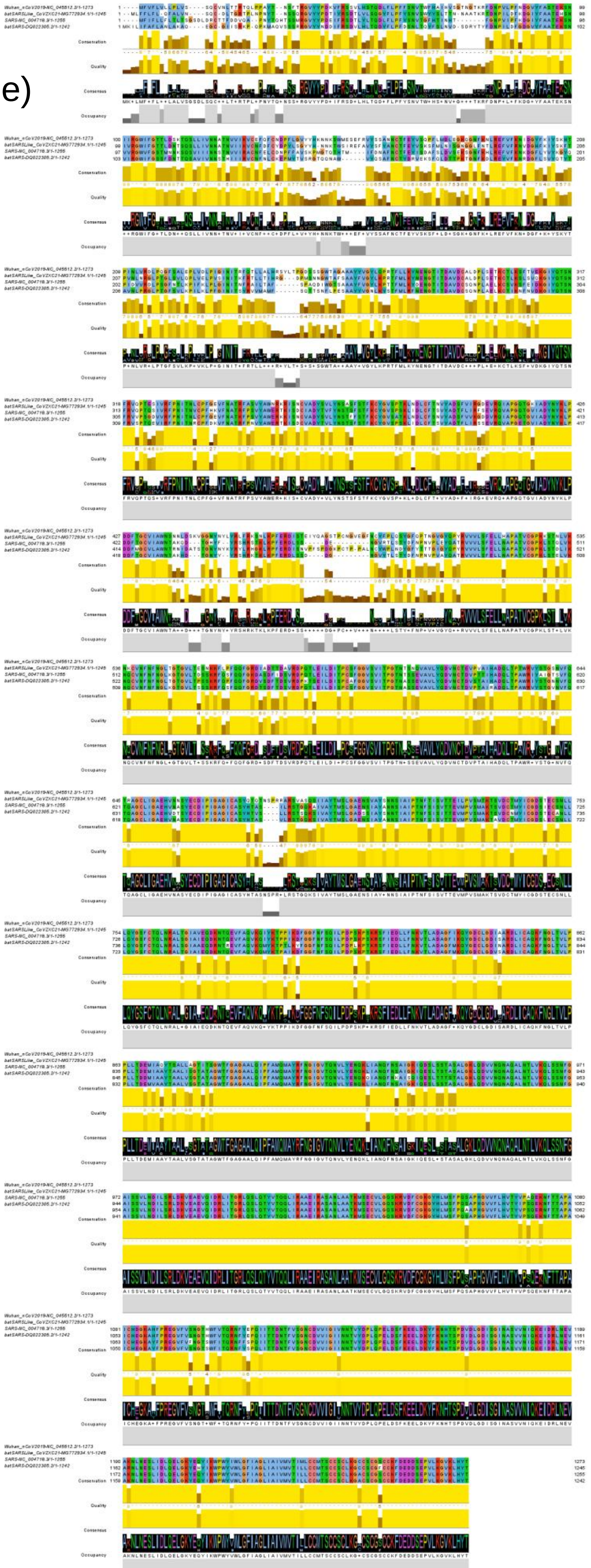
